# Exclusion of PD-1 from the immune synapse: a novel strategy to modulate T cell function

**DOI:** 10.1101/2023.11.16.566907

**Authors:** Luke Yi Hao, Shalom Lerrer, Ruijiang Song, Michael Goeckeritz, Xizi Hu, Adam Mor

**Affiliations:** Columbia Center for Translational Immunology, Columbia University Medical Center, New York, NY 10032; Division of Rheumatology, Department of Medicine, Columbia University Medical Center, New York, NY 10032

**Keywords:** Immune synapse, T cells, monoclonal antibodies, cancer immunotherapy, immune checkpoints

## Abstract

Targeting immune checkpoint receptors on T cells is a common cancer treatment strategy. Frequently, this is accomplished through antibodies targeting the ligand of inhibitory co-receptors. Blocking the immune checkpoint PD-1 binding to its ligands PD-L1 and PD-L2 prevents downstream signaling and enhances anti-tumor T cell responses. This approach improved cancer patients’ outcome. However, only one-third of the patients respond to these treatments. To better understand the mechanism of anti-PD-1 antibodies, we explored the location of PD-1 within the immune synapse. Surprisingly, we discovered that anti-PD-1 antibodies, besides blocking the interaction between PD-1 and its ligands, also removed PD-1 from the synapse. We demonstrated a correlation between removing PD-1 from the synapse by anti-PD-1 antibodies and the extent of T cell activation. Interestingly, a short version of the anti-PD-1 antibody, F(ab’)_2_, failed to remove PD-1 from the synapse and activate T cells. Using syngeneic tumor model, we showed a superior anti-tumor effect to anti-PD-1 antibody over the shorter version of the antibody. Our data indicates that anti-PD-1 antibodies activate T cells by removing PD-1 away from the synapse and changing the location of PD-1 or other immune receptors within immune synapse could serve as an alternative, efficient approach to treat cancer.

## Introduction

Antigen-mediated T cell receptor (TCR) activation depends on the prolonged, stable interaction between a T cell and an antigen-presenting or target cell. The interface between these interacting cells is termed the immunological synapse (IS), and it includes well-organized supramolecular activation clusters (SMACs) [1]. SMACs are concentric rings of segregated proteins sorted by structure and function and driven by the TCR and co-receptor signaling. This structure is essential for proper protein-protein interactions, and the relative proximity between individual proteins within these regions is critical to cellular function. For example, the central SMAC (c-SMAC) is enriched in co-receptors, such as CD4, CD8, CD28, and PD-1; the peripheral SMAC (p-SMAC) is increased in LFA-1/ICAM-1 interaction; and the distal SMAC (d-SMAC) is enriched in F-actin and the transmembrane phosphatase CD45 and CD43 [2].

Due to the tight space between these interacting cells created by integrin-adhesion molecule interactions, the size-based exclusion of molecules from the IS has been previously reported [3, 4]. According to a crystal structure analysis of the IS, the intermembrane distance for receptor-ligand pairs is approximately 15 nm and about 40 nm for integrin-ligand pairs [5, 6]. Using a range of sizes of dextran molecules, it was established that the movement of dextran molecules ≤ 4 nm in and out of the IS was unrestricted. However, the direction of 10 - 13 nm dextran molecules was significantly reduced, and dextran molecules above 32 nm were nearly completely excluded [7]. Further, monoclonal antibodies that correspond to a size of around 15 nm were also excluded from the IS [8].

Targeting immune checkpoints on the surface of T cells is a prevailing approach in cancer immunotherapy [9]. Most immune checkpoint inhibitors that are monoclonal antibodies operate through an assumed mechanism of disrupting receptor-ligand interactions. Antibodies directed toward PD-1, the PD-1 ligand PD-L1, LAG-3, TIM3, TIGIT, and CTLA-4 fall under this category [10]. While these antibodies achieve blockade of receptor-ligand interactions and interfere with outside-in signaling, emerging data suggest this may not be the sole explanation for their efficacy [11]. Further exploring alternative mechanisms to achieve immune checkpoint inhibition promises to expand patient responsiveness and address primary resistance and immune-related adverse events [12].

Because full-length antibodies cannot enter the tight space of the immune synapse, we hypothesized that anti-PD-1 antibodies bind to PD-1 and pull it away from the immune synapse and, therefore, interfere with its function by disrupting the signaling complexes within the synapse. To test this hypothesis, we imaged T cells that express GFP-tagged PD-1 before and after treatment with anti-PD-1 antibodies and recorded PD-1 location and function through in vitro and in vivo systems.

## Results

### PD-1 is localized to the immune synapse

To gain better insight into the location of PD-1 in the immune synapse, we overexpressed GFP-PD-1 in Jurkat T cells and co-cultured these cells with SEE-loaded Raji B cells that expressed mCherry PD-L1. As reported previously, PD-1 was homogenously distributed around the cells during the resting state (Fig. 1A) but localized to the immune synapse in the activated cells (Fig. 1B).

**Figure 1.**
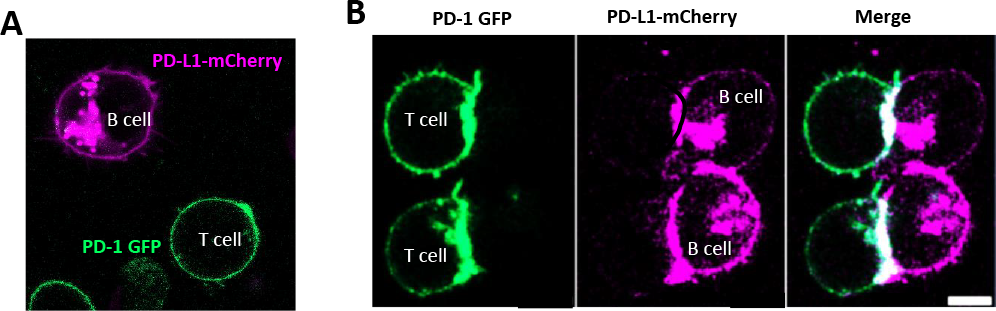
PD-1 is localized to the immune synapse. Jurkat T cells overexpressing GFP-PD-1 co-cultured at rest (A) and with SEE-loaded Raji B cells that expressed mCherry PD-L1 (B). Cells were imaged as being alive; representative cells were shown.

### Full-length anti-PD-1 antibodies removed PD-1 from the immune synapse

It has been reported that antibodies cannot enter the tight immune synapse due to their size [13]. Indeed, we reported that anti-PAG [14, 15] and anti-SLAMF-6 [16, 17] antibodies, due to their size, cannot populate the immune synapse. Thus, we hypothesized that PD-1 is removed from the immune synapse during conjugate formation in the setting of anti-PD-1 antibodies. First, we treated Nivolumab (anti-PD-1 antibody) with pepsin to generate F(ab’)_2_ (Fig. 2Ai). Next, we used flow cytometry to show that Nivolumab F(ab’)_2_ can bind to T cells expressing PD-1 with the same affinity as a full-length Nivolumab (Fig. 2Aii). To better control the experiment, we also treated with Durvalumab, an anti-PD-L1 antibody, with pepsin to generate F(ab’)_2_ (Fig. 2Bi) and showed that its affinity to PD-L1 was maintained (Fig. 2Bii). Next, we treated GFP-PD-1 expressing Jurkat T cells and SEE-loaded Raji B cells expressing mCherry PD-L1 with Nivolumab and its F(ab’)_2_ (Fig. 2Ci) or with Durvalumab and its F(ab’)_2_ (Fig. 2Cii). We quantified the number of cells where PD-1 was recruited to the immune synapse at different concentrations of both antibodies. As shown, while Nivolumab could remove PD-1 from the synapse in most of the cells, its F(ab’)_2_ failed to do that (Fig. 2Di). A similar pattern was observed for Durvalumab and its F(ab’)_2_ (Fig. 2Dii). Altogether, this data suggested that an anti-PD-1 antibody, besides blocking its binding to PD-L1, can also remove its target from the immunological synapse, mediated through its size, without changing binding affinity.

**Figure 2.**
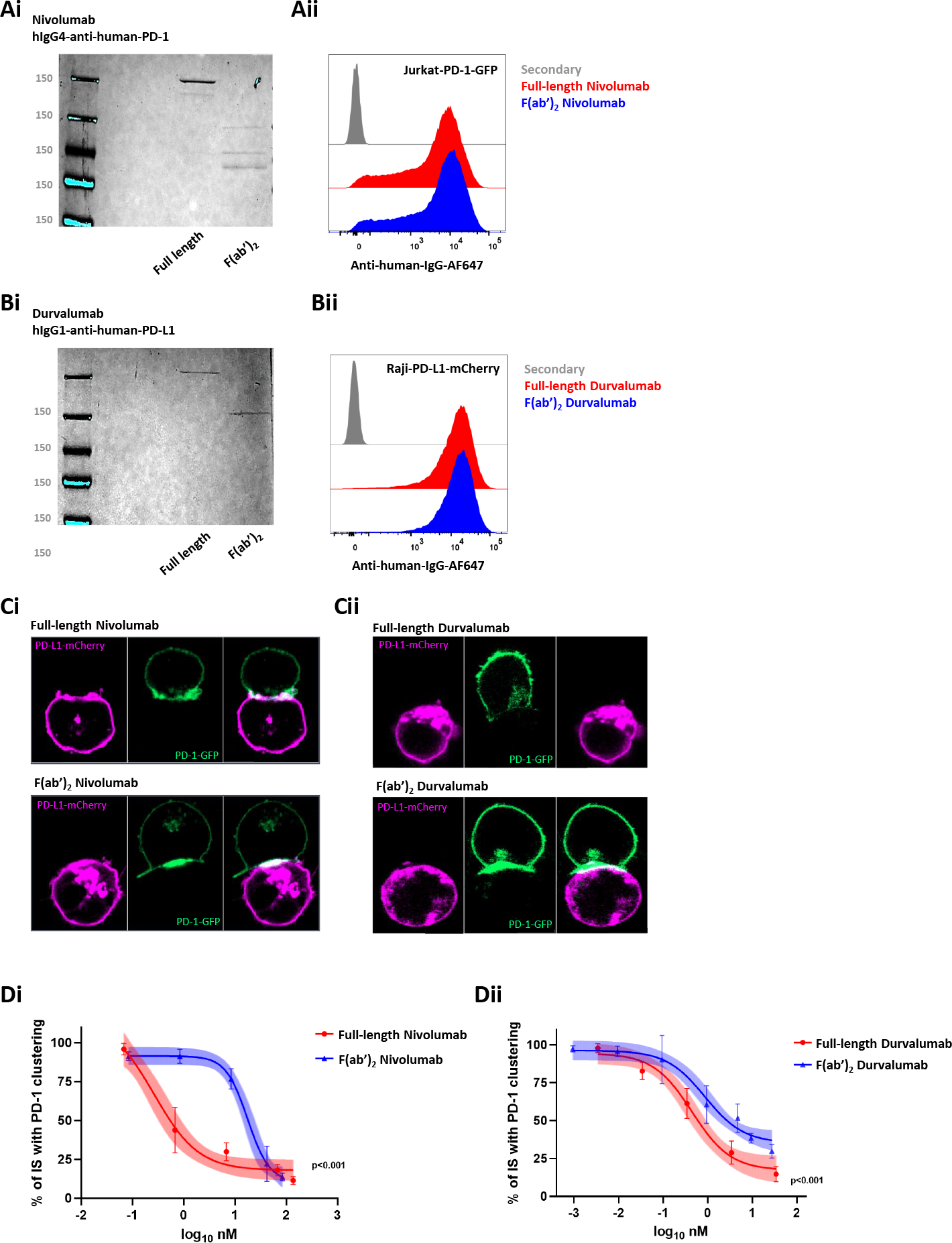
Full-length anti-PD-1 antibodies removed PD-1 from the immune synapse. Coomassie staining of full-length Nivolumab and Nivolumab treated with pepsin (Ai). A representative flow cytometry graph of Jurkat T cells treated with full-length Nivolumab and Nivolumab F(ab’)_2_, as indicated (Aii). Coomassie staining of full-length Durvalumab and Durvalumab treated with pepsin (Bi). Flow cytometry of Raji B cells treated with full-length Durvalumab and Durvalumab F(ab’)_2_, as indicated (Bii). GFP-PD-1 expressing Jurkat T cells co-cultured with SEE-loaded Raji B cells expressing mCherry PD-L1 with Nivolumab and its F(ab’)_2_ (Ci) or with Durvalumab and its F(ab’)_2_ (2Cii) and quantifying the number of cells where PD-1 was recruited to the immune synapse at different concentrations of both antibodies (Di and Dii). N=3, unpaired *t*-test with Welch’s correction, p < 0.001.

### Removing PD-1 from the immune synapse is associated with increased T cell activation

To uncover the biological significance of removing PD-1 from the synapse, we treated Jurkat T cell and Raji B cell with different concentrations of either full-length Nivolumab or its F(ab’)_2_ in the context of SEE (Fig. 3A). The culture media was collected after overnight treatment, and ELISA measured IL-2 levels. As shown, unlike full-length Nivolumab, Nivolumab F(ab’)_2_ could not effectively increase the levels of secreted IL-2. Next, we repeated the same procedure with human PBMCs. As shown, only full-length Nivolumab treatment resulted in increased levels of IL-2 (Fig. 3B) and interferon-gamma (Fig. 3B). To uncover the molecular mechanism and to link our findings of PD-1 signaling, we treated the cells with either full-length Nivolumab or Nivolumab F(ab’)_2_ for 5 minutes, precipitated PD-1 with anti-GFP antibodies and blotted the precipitate with anti-4G10 antibodies to quantify the levels of PD-1 endo domain tyrosine phosphorylation. While Nivolumab F(ab’)_2_ reduced the levels of PD-1 phosphorylation to baseline, the full-length Nivolumab reduced PD-1 phosphorylation levels even further (Fig. 3C), suggesting that tonic signaling downstream of PD-1 can be inhibited not exclusively by interfering with ligand binding, but by physically removing it away from the synapse.

**Figure 3.**
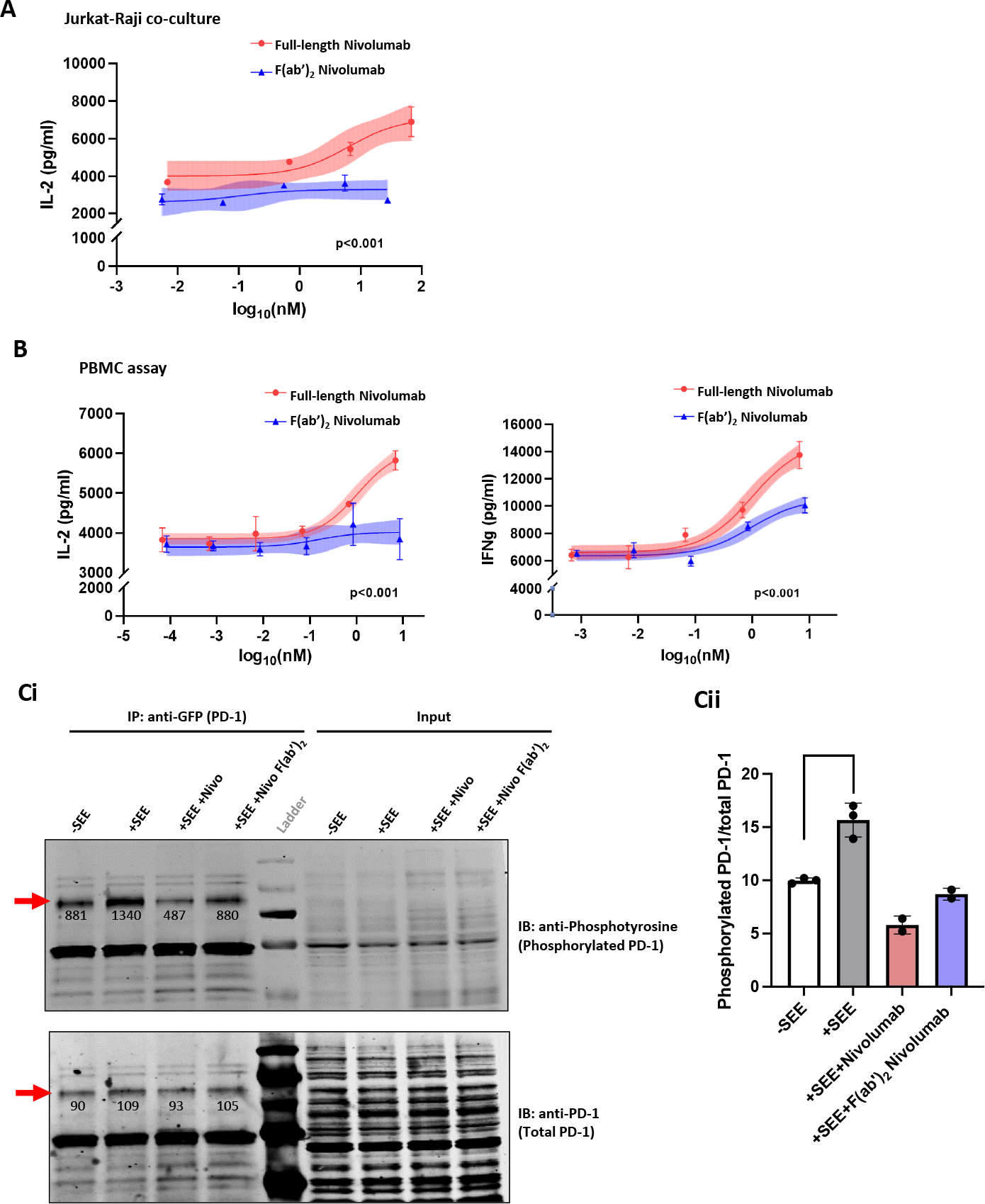
Removing PD-1 from the immune synapse is associated with increased T cell activation. IL-2 levels of Jurkat T cell and Raji B cell with different concentrations of either full-length Nivolumab or its F(ab’)_2_ in SEE (A) context. IL-2 levels, n=3, unpaired *t*-test with Welch’s correction, p < 0.001 and interferon-gamma, n=3, unpaired *t*-test with Welch’s correction, p < 0.001 (B) of human PBMCs assay of cells treated with full-length Nivolumab and its F(ab’)_2_ as indicated and measured by ELISA. The co-immunoprecipitation experiment of Jurkat T cell tread was shown for 5 minutes, followed by PD-1 precipitated with anti-GFP and blotted with anti-4G10 (Ci). Quantitation of the same expedient (Cii). N=3, two-way ANOVA, * p < 0.05.

### Anti-PD-1 F(ab’)_2_ cannot inhibit tumor growth in vivo

To test the role of the anti-PD-1 antibody and its F(ab’)_2_ in vivo, we treated the anti-murine PD-1 antibody with pepsin (Fig. 4A) and confirmed its affinity using an EL4 murine T cell line (Fig. 4B). Next, we used MC38 syngeneic tumor model to test the ability of the antibodies to reduce tumor growth. MC38 inoculated mice were treated with full-length anti-PD-1 antibodies twice a week for four doses or with the same molar amount of anti-murine PD-1 F(ab’)_2_. As shown, the tumors grew faster when the short version of anti-PD-1 was used (Fig. 4C). H&E staining of the tumors at the end of the experiment confirmed an increased number of tumors infiltrating CD3-positive T cells in the mice treated with full-length antibodies (Fig. 4D). Remarkably, both the full-length and the F(ab’)_2_ versions of the anti-PD-1 antibodies were detected on T cells isolated from the spleens of these mice two weeks after the intraperitoneal administration (Fig. 4E).

**Figure 4.**
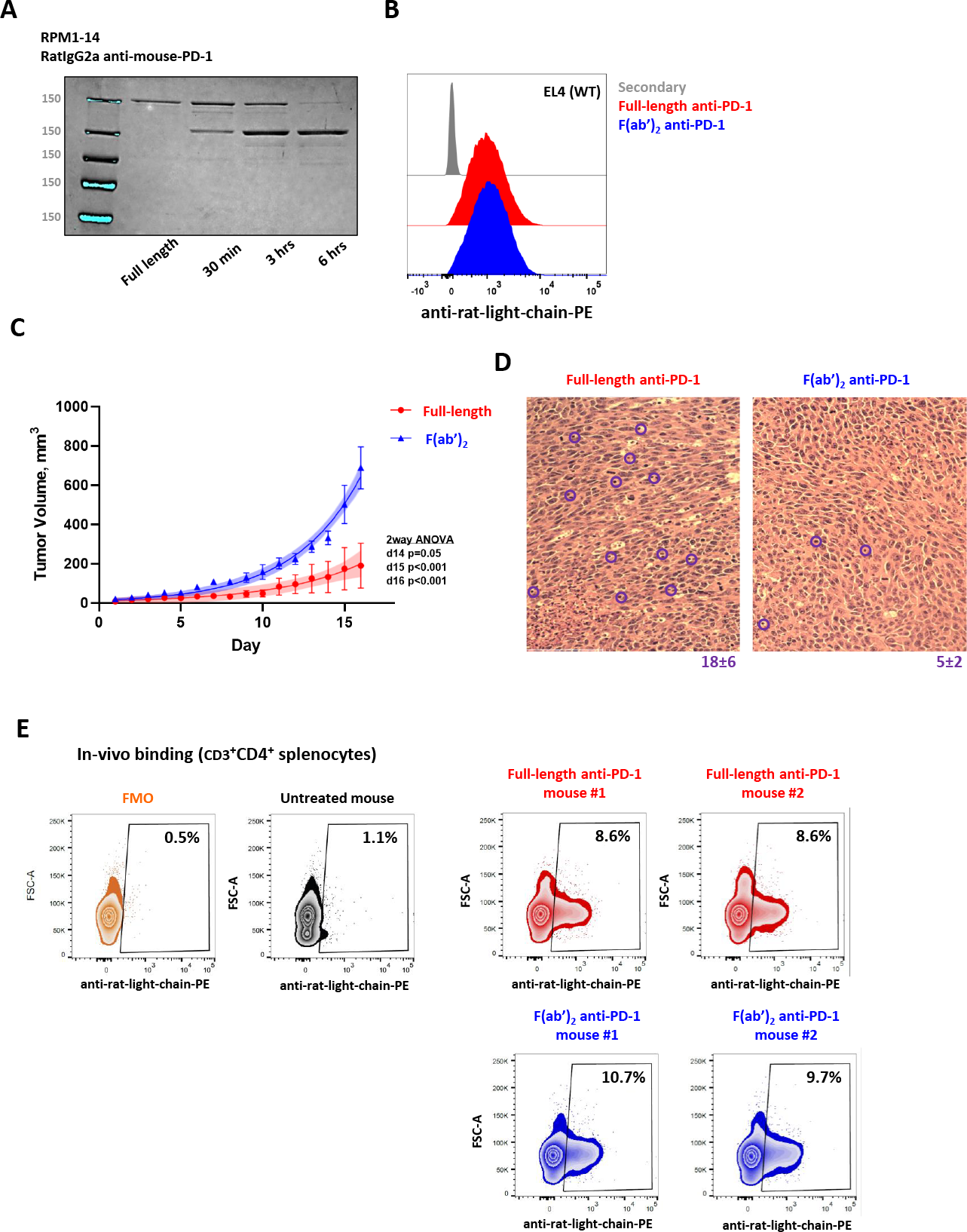
Anti-PD-1 F(ab’)_2_ cannot inhibit tumor growth in vivo. Coomassie staining of full-length anti-murine PD-1 treated with pepsin (Ai). Flow cytometry of EL4 murine T cell line treated with the antibodies, as indicated (B). MC38 syngeneic tumor model showing tumor growth of mice treated with full-length anti-PD-1 antibodies twice a week for four doses or with the same molar amount of anti-murine PD-1 F(ab’)_2_, N=5, two-way ANOVA (C). H&E staining of the tumors at the end of the experiment (D). Flow cytometry of both the full length and the F(ab’)_2_ versions of the anti-PD-1 antibodies using T cells isolated from the spleens of these mice two weeks after the intraperitoneal administration (E). The numbers displayed in the boxes are the percentage of the cells bound with the target full length and the F(ab’)_2_ versions of the anti-PD-1 antibodies.

## Discussion

The PD-1 pathway provides a clear example of how changing a protein’s localization on the cell surface can impact T cell function. During T cell activation, the TCR localizes within micro-clusters within the IS along with the PD-1 [18]. It is well established that PD-1 interacts with Src Homology 2 (SH2) domain-containing proteins upon ligation through phosphorylated tyrosines within its intracellular tail. More specifically, two tyrosine motifs, an immunoreceptor tyrosine-based inhibitory motif and an immunoreceptor tyrosine-based switch motif, are contained within the cytoplasmic tail of PD-1 and become phosphorylated following PD-1 ligation [19]. If PD-1 were re-localized away from the TCR, then the proximity of its cell surface and intracellular binding partners and the TCR would also be disrupted. As a result, these proximity-dependent inhibitors will be unable to act on the TCR cascade, and T cell activation will be enhanced. Perhaps the most well-studied PD-1-interacting protein is the SH2 domain-containing tyrosine phosphatase 2 (SHP2), which binds to the phosphorylated cytoplasmic tail of PD-1. We have previously demonstrated that the ITSM and ITIM contribute to SHP2 localization and full activation [20, 21]. Once activated, SHP2 dephosphorylates critical tyrosine signaling mediators within the CD3 complex and CD28, ZAP70, PI3K, AKT, C3G, and ERK [22, 23]. By removing PD-1 from the TCR microclusters, SHP2 will be mis-localized and remain inactive.

We have shown here that anti-PD-1 monoclonal antibodies block PD-1 interaction with PD-L1 by changing the location of PD-1 away from the p-SMAC toward the d-SMAC. Mechanistically, this is explained by the tight and narrow space between the T cell and the tumor cells, which cannot accommodate the bulky Fc portion of most therapeutic antibodies. Previously, another group showed anti-PD-1 F(ab’)_2_ conjugated PEG-PLGA nanoparticles enhanced immune checkpoint therapy in the same MC38 syngeneic mouse tumor model [24]. We suspected their average bulky size of 260 nm significantly affected the anti-tumor efficacy. To further support that, we were able to show a correlation between the degree of PD-1 exclusion from the IS and its lack of ability to prevent cell proliferation and cytokine secretion. It has been suggested that resistance to PD-1 blockade, at least partially, is secondary to incomplete hindering of PD-1. We recommend addressing this using anti-PD-1 antibodies to remove PD-1 from the IS better than others. Another advantage of this approach is its ability to prevent PD-1 ligand-independent downstream tonic signaling [25]. Since the ability to remove PD-1 from the synapse is not the same with all antibody isotypes, this property should be considered for developing the next-generation anti-PD-1 therapeutics.

Immunotherapies have changed the way we treat patients with cancers. Most of the targets of these therapies are co-receptors expressed on the surface of T cells that are members of the IS [26]. The synapse is a dynamic platform that can initiate downstream signaling by two mechanisms. The first is enabling receptors on T cells to bind to ligands on the target cells, promoting outside-in signaling, or “vertical signaling” [27]. The other mechanism is by forming signaling clusters between proteins located within the membrane of the T cells, a process conceptualized as “horizontal signaling” [28, 29]. The formation of such groups can lead to ligand-independent downstream signaling that could either stimulate or inhibit T cell functions. For example, antibodies to PD-1 block ligand binding and outside-in signaling. However, we have demonstrated that PD-1 inhibition by antibodies may go beyond ligand neutralization, and additional inhibition of this pathway could be accomplished by physically removing PD-1 from the synapse, which also interferes with its horizontal signaling. Similarly, we have demonstrated that moving the adaptor protein PAG away from the IS enhances T cell activation and diminishes PD-1 signaling.

Here, we propose a new design of antibodies targeting proteins in the IS with bulkier sizes, hoping to increase their effectiveness [30]. However, future work is needed to determine the upper limit of the hydrodynamic size of the antibody. In this paper, we have proved that the physical property of an antibody can impact its effectiveness. In the future, other physical properties, such as binding affinity, hinge flexibility, and electrostatic charge, can also be explored.

## Materials and Methods

### General reagents

RPMI 1640, DMEM, Dulbecco’s PBS (DPBS), penicillin/streptomycin, and fetal bovine serum (FBS) were purchased from Life Technologies. Lymphoprep was purchased from StemCell Technologies. Staphylococcal Enterotoxin E (SEE) was purchased from Toxin Technology (ET404).

### Cell culture, stimulation

Jurkat T cells and Raji B cells were obtained from the American Type Culture Collection (TIB-152 and CCL-86). These cells were maintained in 5% CO_2_ at 37°C in RPMI 1640 supplemented with 10% FBS and 1% penicillin/streptomycin. Jurkat T cells and Raji cells were stably expressing pHR-PD-1-GFP vector and pHR-PD-L1-mCherry vector, respectively, as described previously [31]. The imaging experiments combined Jurkat T cells and Raji B cells with a 1:1 ratio. For immunoprecipitation for phosphorylated PD-1 level, Jurkat T cells and Raji B cells were mixed with a 3:1 ratio.

### Imaging and microscopy

Raji B cells (1 million cells/mL) loaded with 0.1 ug/mL SEE were cocultured with Jurkat T cells (1 million cells/mL) with a 1:1 ratio before being treated with the Nivolumab and Durvalumab antibodies. For conjugate experiments, live cell co-cultures were imaged for 30 minutes using confocal microscopy (Zeiss LSM 900). IS and PD-1 clustering were manually counted in ZEN 3.3 (blue edition) for quantification.

### Antibody processing and validation

Clinical grade Nivolumab fragments, clinical grade Durvalumab fragments, and rat-anti-mouse-PD-1 antibody (catalog no. BE0146; InVivoMAb) were prepared by Pierce F(ab’)_2_ Macro Preparation Kit (catalog no. 44988). The sizes of antibody fragments were measured through Tris-glycine PAGE, stained with Coomassie Brilliant blue before destain, and visualized on a gel reader. Intensity was measured by Image Studio Ver 5.2.

### Western blotting

After running the previous SDS-PAGE gel for an ideal time, proteins were transferred for 30 min at 25V, the nitrocellulose membrane was blocked with 5% BSA in TBS containing 0.05% Tween 20 and blotted overnight with primary antibody prepared in TBST containing 2.5% BSA. The membrane was developed using a secondary fluorescent antibody and acquired on an Odyssey CLx Imaging system.

### Immunoprecipitation

Raji B cells over-expressing PD-L1 and Jurkat T cells over-expressing PD-1 were lysed in cold IP lysis buffer (25mM Tris-HCL, 150mM NaCl, 1mM EDTA, 0,5% NP-40, 5% Glycerol). The lysis process was carried out at 4°C for 30 minutes. The lysates were centrifuged for 10 minutes at 12,000 g and 4°C. Lysates were then used to immunoprecipitate PD-1-GFP using anti-GFP antibody-conjugated agarose beads (MBL) according to the manufacturer’s protocol. The PD-1-GFP protein was then separated from the beads by sample buffer (2x Laemmli buffer, boiled at 95°C for 10 minutes) and loaded onto SDS-PAGE gel.

### IL-2 and IFN-gamma ELISA

To determine the concentration of secreted cytokines following different stimulation conditions, the human IL-2 ELISA kit (catalog no. 431801; BioLegend) and human IFN-gamma kit (catalog no. 430101; BioLegend) were used according to the manufacturer protocols.

### Flow cytometry

The binding of full-length and fragmented Nivolumab was tested using Jurket-PD-1-GFP cells, the binding of full-length and fragmented Durvalumab was tested using Raji-PD-L1-mcherry cells, and the binding of full-length and fragmented anti-mouse-PD-1 was tested using wild type EL4 cells. In all cases, 0.5 million cells/test were incubated with equal molar concentrations of full-length/fragmented antibodies in FACS buffer (2% FBS in PBS) for 30 minutes at 4°C. Following 2 washes, the cells were stained for 30 minutes at 4°C with a fluorescently labeled secondary antibody – anti-human-IgG-F(ab’)_2_ (Jackson ImmunoResearch #109-606-097, used at 1:400) for Nivolumab and Durvalumab-stained samples, and anti-rat-Ig-light-chain (Biolegend #407805, used at 1:250) for anti-mouse-PD-1-stained samples, washed twice, and acquired using BD LSRII flow cytometer. Data was analyzed using FlowJo software (version 10.7.1).

### Mice and tumor cell lines

The Columbia University Institutional Animal Care approved the Animal Studies and Use Committee (IACUC). Female, 8-week-old C57BL/6 mice were used. The murine colon adenocarcinoma (MC38) colon carcinoma cells were purchased from Kerafast (catalog no. ENH204-FP). The MC38 cells were maintained in DMEM supplemented with heat-inactivated fetal bovine serum (10%) and penicillin/streptomycin (1% 10,000 U/mL stock) and grown at 37°C with 5% CO_2_. Cells were passaged before storage and thawed and passaged twice before implantation for all described tumor experiments. All cell lines were determined to be free of mycoplasma (Lonza).

### Tumor model

MC38 (200x10^5^) cells were implanted subcutaneously in the right hind flank of 8-week-old mice. Tumor growth was monitored using electronic calipers and calculated using V = length*width^2^*0.52. When tumor volume reached 50 mm^3^, the mice started receiving treatments (2 doses per week, four doses total). Tumor sizes were tracked along the course. Mice were eventually sacrificed, and tumors were collected for histology.

### Tumor-infiltrating histology

For H&E, tumors were fixed in 10% neutral buffered formalin, and then paraffin was embedded and cut into five μm sections. Slices were stained with H&E. T cells were qualified by counting mononuclear cells in a high-power field.

### Statistics

Values are reported as mean ± SEM. Statistical graphs were analyzed on GraphPad Prism 9, using an unpaired *t*-test with Welch’s correction and ANOVA two-way test.

### Ethics

The Institutional Review Board at Columbia University Medical Center approved the study, and all donors provided informed consent.

## Data Availability Statement

All data are available upon reasonable request from the corresponding author, Adam Mor.

## Acknowledgment

This work was supported by grants from the NIH (AI125640, CA231277, AI150597, AI175498).

## Author Contributions

L.Y.H. performed the experiments, analyzed the data, and drafted the manuscript. S.L. helped improve the process of the experiments and analyze the data. R.S., M.G., and X.H. helped improved the process of the experiments. A.M. sketched out the experiments, revised the manuscript, provided material support, and offered reagents and data analysis tools. The manuscript was read, verified, and approved by all authors.

## Declaration of Interests

The authors declare no competing interests.

## Notes

### Competing Interest Statement

The authors have declared no competing interest.

